# A more accurate method for colocalisation analysis allowing for multiple causal variants

**DOI:** 10.1101/2021.02.23.432421

**Authors:** Chris Wallace

## Abstract

In genome-wide association studies (GWAS) it is now common to search for, and find, multiple causal variants located in close proximity. It has also become standard to ask whether different traits share the same causal variants, but one of the popular methods to answer this question, coloc, makes the simplifying assumption that only a single causal variant exists for any given trait in any genomic region. Here, we examine the potential of the recently proposed Sum of Single Effects (SuSiE) regression framework, which can be used for fine-mapping genetic signals, for use with coloc. SuSiE is a novel approach that allows evidence for association at multiple causal variants to be evaluated simultaneously, whilst separating the statistical support for each variant conditional on the causal signal being considered. We show this results in more accurate coloc inference than other proposals to adapt coloc for multiple causal variants based on conditioning. We therefore recommend that coloc be used in combination with SuSiE to optimise accuracy of colocalisation analyses when multiple causal variants exist.

## Introduction

Colocalisation is a technique used for assessing whether two traits share a causal variant in a region of the genome, typically limited by LD. In its original form, it made the simplifying assumption that the region harboured at most one causal variant per trait[1]. The approach proceeds by enumerating all *variant-level hypotheses* - the possible pairs of causal variants (or none) for the two traits - and the relative support for each in terms of Bayes factors. Each one of these combinations is associated to exactly one *global hypothesis*

*H*_0_ : no association with either trait in the region

*H*_1_ : association with trait 1 only

*H*_2_ : association with trait 2 only

*H*_3_ : both traits are associated, but have different single causal variants

*H*_4_ : both traits are associated and share the same single causal variant

The Bayes factors for each of these global hypotheses may be calculated by summing the related Bayes factors for each variant-level hypothesis, and simple combination with prior probabilities of each hypothesis allows us to calculate posterior probabilities.

This simple summation is enabled by the single causal variant assumption, which implies that each pair of variants being causal for the two traits are mutually exclusive events. However, the assumption is unrealistic, as multiple causal variants may exist in proximity, which also challenges the definition of colocalisation as presented above as none of the global hypotheses encompass multiple causal variants.

In previous work,[2] we allowed for multiple colocalisation comparisons to be performed in a region, each labelled by a pair of SNPs tagging each of the distinct causal variants for each trait. Thus, if trait 1 had two causal variants tagged by SNPs A and B and trait 2 had one, tagged by SNP C, we would conduct two colocalisation analyses, to ask whether A and C corresponded to a shared causal variant, and whether B and C corresponded to a shared causal variant. This allows the simple combination of Bayes factors through summation, but explicitly assumes that data can be decomposed into layers corresponding to the causally distinct signals. The stepwise regression approach upon which conditioning is based is known generally to produce potentially unreliable results[3], a phenomenon that can be exacerbated by the extensive correlation between genetic variants caused by linkage disequilibrium (LD)[4]. Thus, this solution remains unsatisfactory.

A suite of Bayesian fine-mapping methods have been developed recently which calculate posterior probabilities of sets of causal variants for a given trait[5, 6, 7]. However, the marginal posterior probabilities calculated from these are no longer mutually exclusive events, so they could not be easily adapted to the colocalisation framework. An alternative would be to consider all possible combinations of models between two traits, but this combinatorial problem is computationally expensive[4]. Recently, the Sum of Single Effects (SuSiE) regression framework[8] was developed which reformulates the multivariate regression and variable selection problem as the sum of individual regressions each representing one causal variant of unknown identity. This allows the distinct signals in a region to be estimated simultaneously, and enables quantification of the strength of evidence for each variant being responsible for that signal. Conditional on the regression being considered, the variant-level hypotheses are again mutually exclusive. Here we describe the adaptation of coloc, allowing for multiple labelled comparisons in a region, to use the SuSiE framework and demonstrate improved efficacy over the previously proposed approaches.

## Methods

### Adaptation of coloc approach

The new `coloc.susie` function in the coloc package (https://github.com/chr1swallace/coloc/tree/susie) takes a pair of summary datasets in the form expected by other coloc functions, runs SuSiE on each and performs colocalisation as described below. We use the `susie_rss()` function in the susieR package to fine-map each summary statistic dataset, run with default options, although the `susie.args` argument in `coloc.susie()` allows arguments to be supplied to `susie_rss()`. SuSiE returns a matrix of variant-level Bayes factors for each modelled signal and a list of signals for which a 95% credible set could be formed, corresponding to a subset of rows in the matrix of Bayes factors. These rows are then analysed in the standard coloc approach, for every pair of regressions with a detectable signal across traits. Explicitly, if *L*_1_ and *L*_2_ signals are detected (have a credible set returned) for traits 1 and 2 respectively, then the colocalisation algorithm is run *L*_1_ × *L*_2_ times. Thus, the user is presented with a list of tag SNPs per signal for each trait, and the matrix of pairwise posterior probabilities of *H*_4_ may be examined to infer which, if any, pairs of tags represent the same signal.

### Decreasing the computational burden

While SuSiE has been shown to have greater accuracy than other fine-mapping approaches[8], `susie_rss` becomes computationally expensive as the number of variants in a region increases. Both coloc and fine-mapping require dense genotyping data to make an accurate assessment, so computational complexity can become a burden in larger genomic regions. In such regions, we propose that the fine-mapping result can be approximated by running SuSiE only on a subset of variants with some weak evidence for association (excluding those with larger *p* values), and setting the Bayes factors at those variants not considered to the minimum Bayes factor over all other variants. We use the term “trimming” to describe this approach.

### Simulation strategy

We examined the performance of the approximation described above to decrease the computational burden, and of using SuSiE with coloc by simulation. We used lddetect[9] to divide the genome into approximately LD-independent blocks, and extracted haplotypes from the EUR samples in 1000 Genomes phase 3 data, consisting of 1000 contiguous SNPs with MAF > 0.01. We simulated case-control GWAS summary statistics for a study with 10,000 cases and 10,000 controls, corresponding to the LD and MAF calculated from these haplotypes using simGWAS[10], with one or two common causal variants (MAF > 0.05) chosen at random and log odds ratios sampled from *N* (0, 0.2^2^). We discarded any datasets which did not have a minimum *p* < 10^−6^ to match our expectation that fine-mapping and colocalisation are only conducted when there is at least a nominal signal of association. We simulated 100 such datasets for each of 100 randomly selected LD blocks, and sampled from these sets of summary data for all the simulations detailed below.

To investigate the validity of the approximation, we repeatedly simulated GWAS summary data for a single trait with one or two causal variants in small or large genomic regions (1000 or 3000 SNPs, where 3000 SNP regions were constructed by concatenating three 1000 SNP datasets). We simulated 2000 such examples, and analysed each dataset once using all SNPs and three times, trimming SNPs with |*Z*| <0.5, 1.0, and 1.5. To derive a summary measure of performance, we considered the credible sets identified as individual signals, and calculated the total posterior inclusion probability (PIP) at each SNP as the sum of PIP across these signals. The number of credible sets may change between trimmed and untrimmed datasets, either increasing or decreasing. We therefore derived two final summary measures as the difference between sum of this total PIP at the simulated causal variants in trimmed and untrimmed data, and the difference between the sum of total PIP across all the other (non-causal variants) between trimmed and untrimmed data. If trimming is a successful approximation, these differences should both be 0. If trimming decreases accuracy of fine-mapping, we might see the sum of PIP assigned to causal variants decrease, whilst if it introduces false positive signals, we might see the sum of PIP assigned to non-causal variants increase.

For colocalisation, we simulated data for two traits in the same way, such that each trait had one or two causal variants and each pair of traits shared zero, one or two causal variants. We simulated 10,000 examples from each collection, with each example analysed independently. Analysis compared different approaches:

1. single causal variant coloc analysis of every pair of traits
2. multiple causal variant coloc analysis using a conditioning approach to allow for multiple causal variants, iterative mode
3. multiple causal variant coloc analysis using a conditioning approach to allow for multiple causal variants, “all but one” mode
4. multiple causal variant coloc analysis using SuSiE to allow for multiple causal variants, including data trimming based on |*Z*| score

Conditioning can be run in two modes. Assume that stepwise regression detects two signals, tagged by SNPs A and B. In the iterative mode, we first use the raw data in a first step, and then the data conditioned on A in a second step. This corresponds to how stepwise identification of independent signals in GWAS is commonly approached. An alternative is to condition on B in the first step, and A in the second step, attempting to isolate the separate signals. This corresponds more closely to the hope in multiple causal variant coloc that we can decompose the data into layers corresponding to the separate signals. However, because the identification of the second signal B is likely to be more uncertain than A (because it is weaker, and was detected through conditioning on the already uncertain A), it may introduce further error.

In order to assess the accuracy of each coloc analysis, we needed to assess whether the comparison corresponded to a case of shared or distinct causal variants. For each signal passed to coloc, we identified the variant with the highest posterior probability of causality, *v*_1_ and *v*_2_ for traits 1 and 2 respectively (it is possible that *v*_1_ = *v*_2_). We then labelled the variant *v*_*i*_ (*i* = 1, 2) according to the rules:

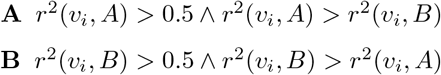

– otherwise

If either of the variants was labelled “-” then the comparison was labelled “unknown”. Otherwise it was labelled by the concatenation of the two labels. We compared the average posterior probability profiles between methods, stratified according to this labelling scheme.

## Results

First we assessed the impact of trimming data on the accuracy and speed of SuSiE. We found that trimming had a very small effect on PIP estimates at the causal variants (Fig 1). Interestingly, when estimates did change, they were more likely to detect a true signal after trimming than lose a true signal (approximately 1-2% of simulations related led to true causal variants that were discovered only after trimming, while ≤ 1% of simulations led to true causal variants being discovered in the full data but not in the trimmed data). False signals were also more likely to be detected after trimming, however. This was more extreme with larger |*Z*| thresholds and in simulations with two rather than one causal variants, when over 2% of simulations resulted in signals being detected at non-causal variants after trimming at |*Z*| < 1.5. One might expect the situation to worsen if the number of causal variants increased. Thus, trimming is expected to introduce false positives at a higher rate than it might increase detection of true positives, although it did reduce the median time for a SuSiE run per region more than ten fold (Fig 2).

**Figure 1:**
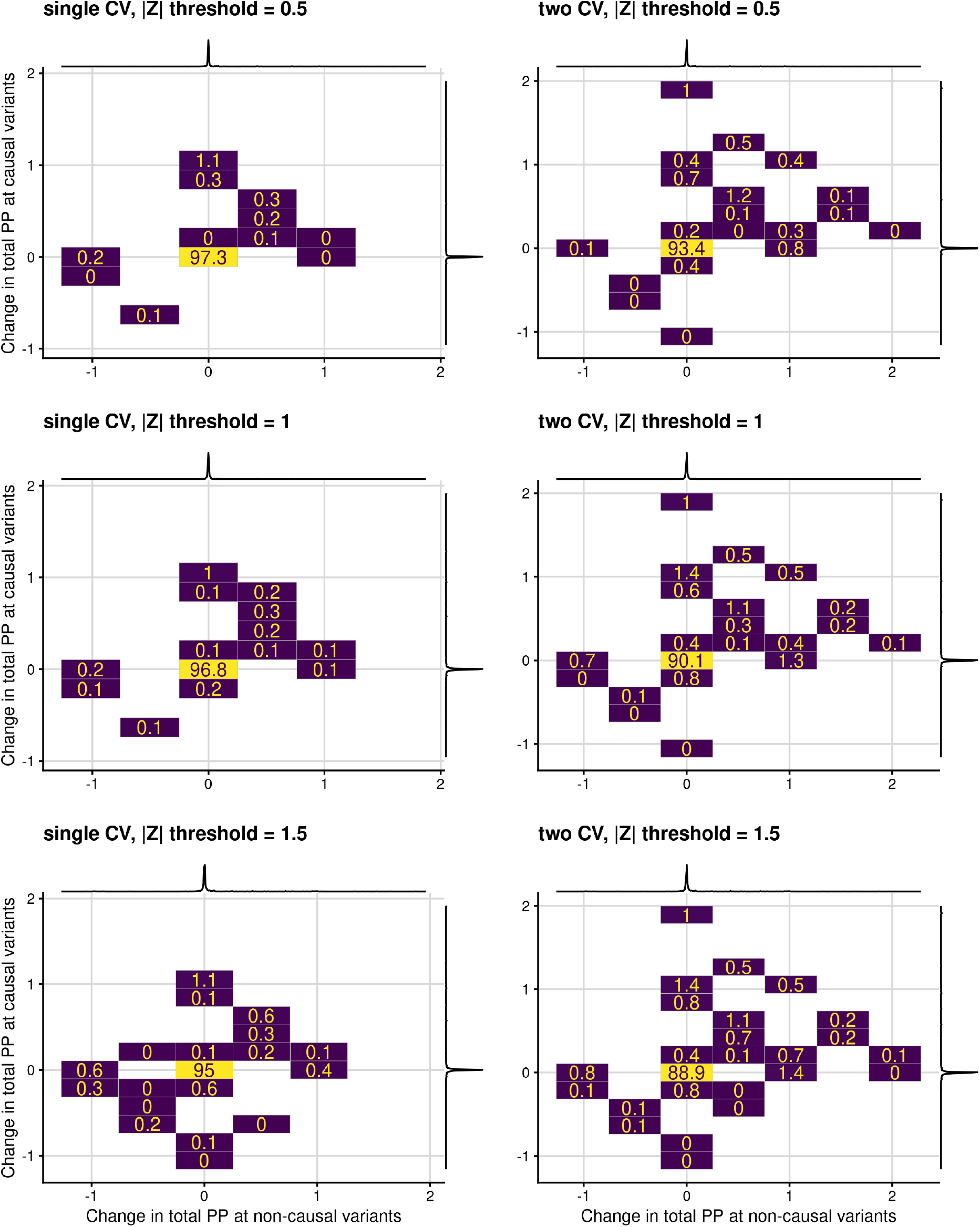
Difference in the sum of estimated PIP at the causal variant(s) (y-axis) versus non-causal variants (x-axis) between analysis with the full model and data trimmed to |*Z*| above some threshold. The percentage of 2000 simulations falling in each region is shown to the nearest 1 decimal place (note that “0” indicates < 0.05%). Marginal densities show the concentration of observations in either direction around (0, 0). Datasets all had 1000 SNPs, but differ in the number of causal variants (1 or 2) and the |*Z*| threshold used for trimming.

**Figure 2:**
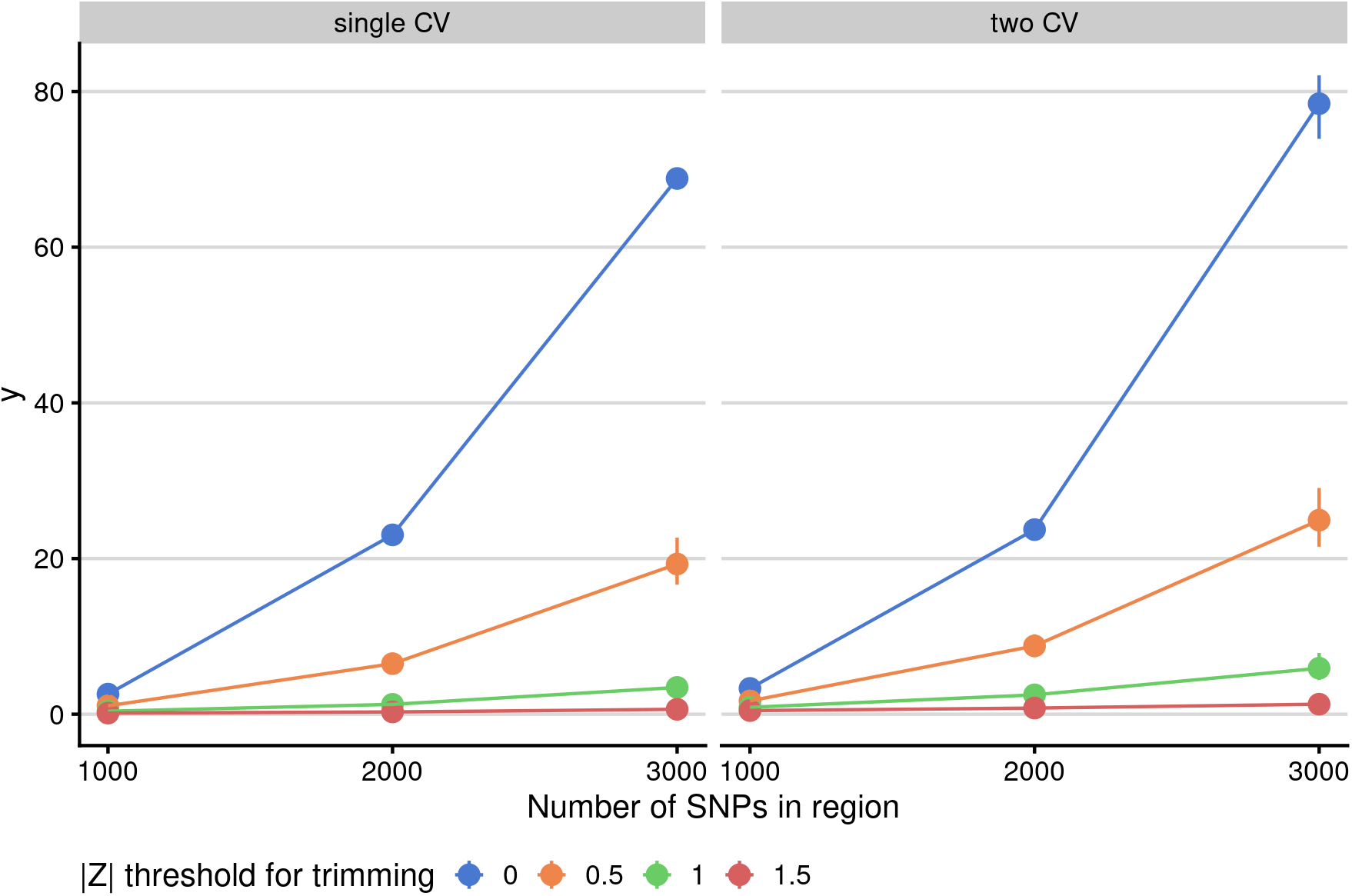
Time to run SuSiE per region in relation to the number of SNPs in the region (1000, 2000 or 3000), the number of causal variants (1 or 2) and the threshold used to trim SNPs by their —Z— scores. The point shows the median time over 1000 simulations, and the vertical range its interquartile range.

The results of the coloc simulation study are given in Supplementary Table 1, and presented graphically in Fig 3 and Supplementary Fig 2. We found that inference with SuSiE coloc was broadly equivalent to that with other approaches when both traits really did contain only a single causal variant (top two rows of Fig 3). When either one or both traits had two causal variants (bottom two rows of Fig 3), all methods apart from single coloc were broadly similar in terms of favouring *H*_4_ when comparing truly colocalising signals (“AA” or “BB” comparisons). Single coloc tended to equivocate between *H*_3_ and *H*_4_ when testing AB-like signals in the presence of a shared causal variant (ie where the peak signals in each trait related to distinct causal variants) which should be inferred *H*_3_. This relates to a known feature of coloc, which may detect the colocalising signal even when additional non-colocalising signals are present,[1].

**Figure 3:**
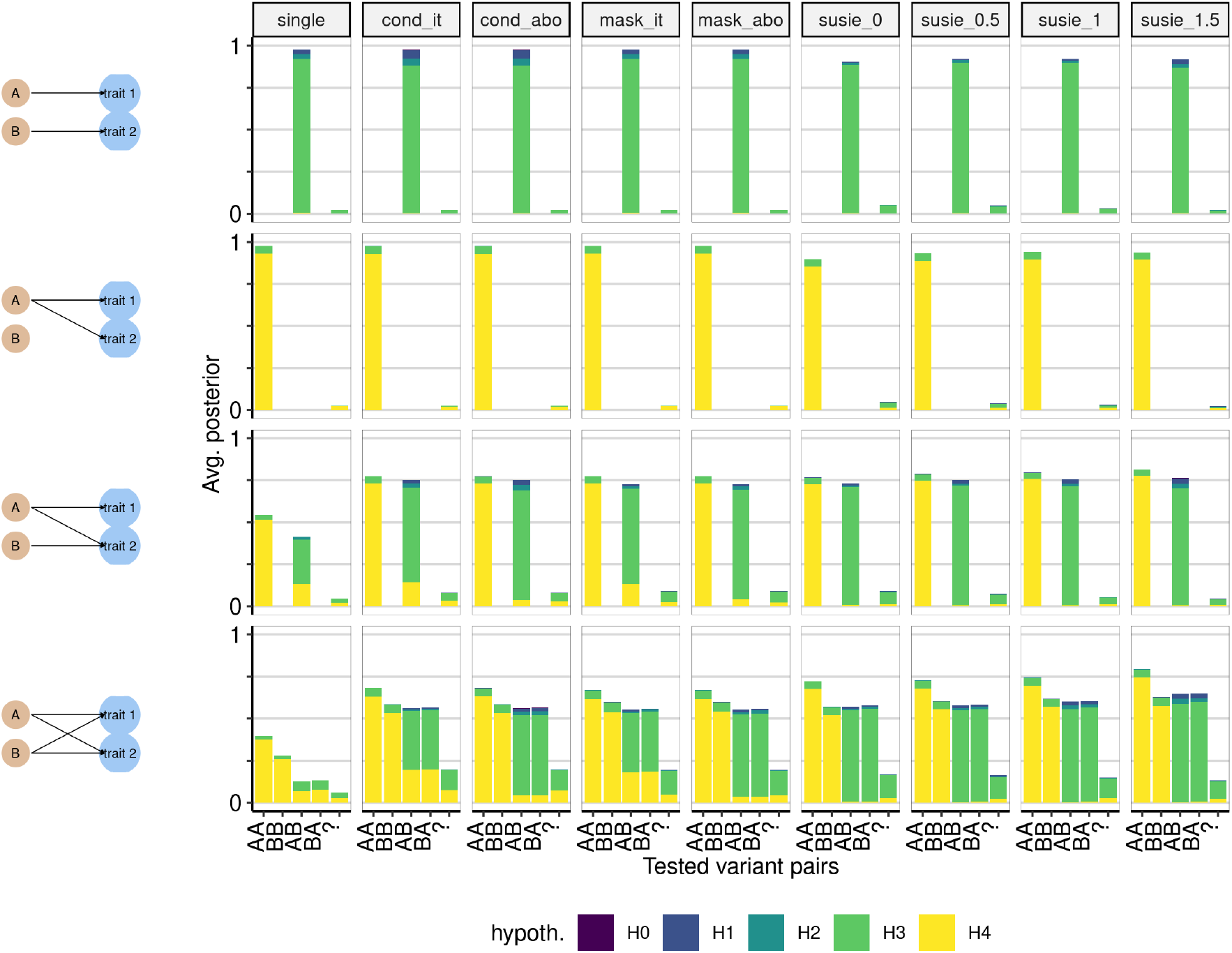
Average posterior probability distributions in simulated data. The four classes of simulated datasets are shown in four rows, with the scenario indicated in the left hand column. For example, the top row shows a scenario where traits 1 and 2 have distinct causal variants A and B. Columns indicate the different analysis methods, with susie x indicating that SuSiE was run with data trimmed at |*Z*| < x, cond it indicating that conditioning was run in iterative mode, and cond abo indicating it was run in “all but one” mode. For each simulation, the number of tests performed is at most 1 for “single”, or equal to the product of the number of signals detected for the other methods. For each test, we estimated which pair of variants were being tested according to the LD between the variant with highest fine-mapping posterior probability of causality for each trait and the true causal variants A and B. If *r*^2^ > 0.5 between the fine-mapped variant and true causal variant A, and *r*^2^ with A was higher than *r*^2^ with B, we labeled the test variant A, and vice versa for B. Where at least one test variant could not be unambigously assigned, we labelled the pair “?”. The total height of each bar represents the proportion of comparisons that were run, out of the number of simulations run, and typically does not reach 1 because there is not always power to perform all possible tests. Note that because we do not limit the number of tests, the height of the bar has the potential to exceed 1, but did not do so in practice. The shaded proportion of each bar corresponds to the average posterior for the indicated hypothesis, defined as the ratio of the sum of posterior probabilities for that hypothesis to the number of simulations performed. Each simulated region contains 1000 SNPs.

This feature also presents problems for the conditioning approach, as demonstrated by the high average posterior probability for *H*_4_ in the “AB” comparisons, one of which is examined in detail in Fig 4. In this example, trait 1 has one causal variant, A, whilst trait 2 has two, A and B, with B having slightly greater significance. In the first round of analysis by the conditioning method, the original sets of summary statistics are passed to coloc. Because A is the stronger effect for trait 1, the test is labelled “AB”, but gives a high posterior to *H*_4_ because there is one shared causal variant (A). Then the stronger effect, B, is conditioned out, and the analysis rerun with trait 1, and trait 2 conditioned on B. This test again gives a high posterior for *H*_4_. This situation is confusing, because the same signal in trait 1 appears to colocalise with different signals in trait 1. SuSiE models both signals simultaneously, so we can attempt to colocalise trait 1 with each signal independently, finding high *H*_3_ for one and high *H*_4_ for the other. If we were confident we could infer both the exact number of independent signals and their identity correctly by conditioning, we could attempt to emulate this in the conditioning, using the “all but one” rather than “iterative” mode. This does result in better average performance than the iterative mode (Fig 3). However it is often outperformed by SuSiE. Supplementary Fig 3 shows an example where the stepwise approach is less able to correctly identify the separate signals. The A signal is not well identified, and therefore not be adequately conditioned out, which may results in two apparently different comparisons with trait 1 which both produce a high *H*_4_. In this example too, SuSiE more correctly produces two comparisons, one with high *H*_3_ and one with high *H*_4_.

**Figure 4:**
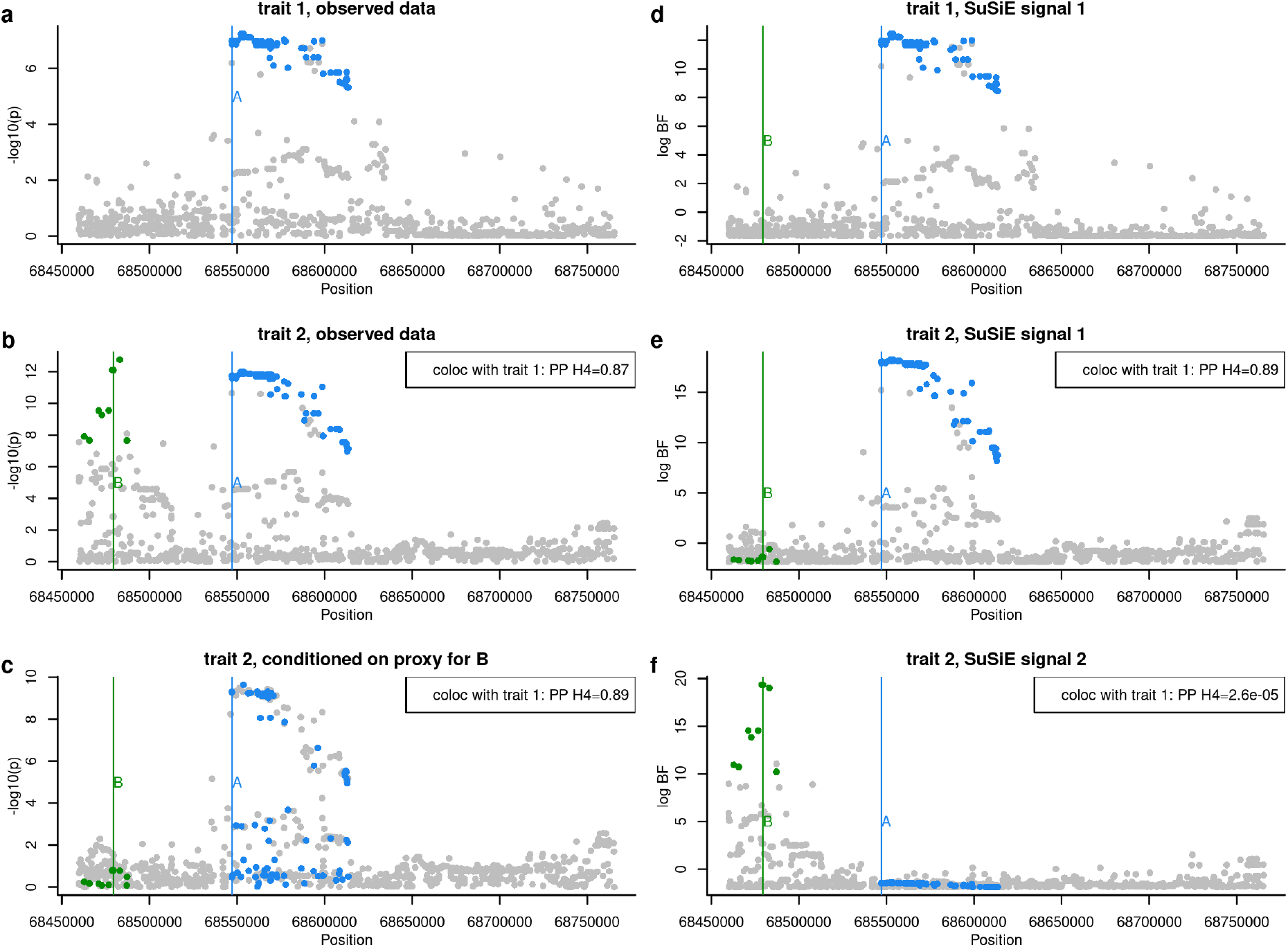
Example where the conditional coloc approach, run in iterative mode, finds misleading results. **a** and **b** show the observed data (-log p values) for traits 1 and 2 respectively. Conditioning identifies a second independent signal for trait 2, and the results of conditioning on the strongest signal is shown in **c**. Coloc comparisons are based on (*a, b*) and (*a, c*) and both find the posterior probability (PP) of *H*_4_ is > 0.8. SuSiE analysis of the same data finds one signal in trait 1, and log Bayes factors (BF) for this signal are shown in **d**. It finds two signals for trait 2, and the log BF for these are shown in **e** and **f**. Coloc comparisons are based on (*d, e*) and (*d, f*) and find PP of *H*_4_ of > 0.9 and < 10^−4^ respectively. Blue and green points are used to highlight SNPs in LD with (*r*^2^ > 0.8) the true causal variants A and B respectively.

Interestingly, the fine-mapping false positives introduced by trimming do not appear to affect the coloc performance. Results are similar across the range of different possible thresholds for trimming, with trimming even producing a slight improvement in coloc accuracy, particularly in regions with more SNPs (Supplementary Fig 2).

## Discussion

While coloc has been a popular method for identifying sharing of causal variants between traits, the common simplifying assumption of a single causal variant has been criticised, because it does not accord with findings that causal variants for the same trait may cluster in location (e.g. because they act via the same gene)[11]. Using the new SuSiE framework to partition the problem into multiple coloc comparisons and assuming the single causal variant assumption holds in each appears to resolve this issue better than the previously proposed conditional approach. It allows multiple signals to be distinguished, and then colocalisation analysis conducted on all possible pairs of signals between the traits.

Despite the adoption of a novel iterative procedure to fit the SuSiE model, the procedure is still slow for large regions with many SNPs, which can be a barrier to its adoption for a technique like coloc which has always boasted speed as an advantage. We note that the package susieR is still being developed. The most computationally expensive step in `susie_rss` is the eigen decomposition of the LD matrix, which is *O*(*p*^3^) where *p* is the number of SNPs. In the case where multiple pairs of traits are being considered for colocalisation, computing this decomposition once in advance could be used instead to improve speed. Alternatively, it may be that with standardised datasets covering the same sets of SNPs, such eigen decompositions could be precomputed and stored. Finally, further development of `susie_rss` may lead to avoiding the eigen decomposition step altogether.

In this manuscript, we considered a simple approach, approximating the SuSiE posterior by using a trimmed set of data, discarding SNPs with |*Z*| scores below some small threshold, on the assumption that a causal SNP with detectable association should produce a *Z* score of reasonable magnitude. (For reference, whilst we only consider discarding SNPs with |*Z*| < at most, the standard genome-wide significance threshold of *p* < 5 × 10−8 corresponds to |*Z*| > 5.45). Thus, this approximation makes the assumption that true causal variants will have at least some weak marginal evidence of association. We note that it is possible to construct examples which will violate this assumption, for example if two causal variants in strong LD but with opposite directions of effects exist. Further, in simulated data here, we found that trimming can increase false positive signals when fine-mapping, a phenomenon previously noted[12]. Thus we suggest that trimming is inappropriate when fine-mapping is the end-goal of any study, especially given that false positives in fine-mapping may result in substantial costs if followed-up in wet-lab experiments. However, these fine-mapping false positives did not appear to increase false positives in coloc, perhaps because the occurence of false positives was relatively rare, such that it did not occur often in both members of a pair of datasets, and/or perhaps because when false positives did occur, they were focused on different SNPs in the two datasets so were unlikely to generate support for *H*_4_, the hypothesis of most interest. Given this, we suggest a threshold of |*Z*| < 1 may be acceptable to allow SuSiE coloc to run at speed in larger regions, but leave the threshold as a user-set parameter which is 0 by default, and which we recommend should be reported along with any results. For the most noteworthy results, we recommend analysis should be repeated without trimming to ensure inference is robust to the approximation.

This manuscript presents one approach to colocalisation in the case of multiple causal variants, that assumes that distinct signals can be decomposed even if physically proximal, which SuSiE appears to do admirably well. This framing of the colocalisation problem implicitly assumes there are a finite number of causal variants for any trait which can be identified, and that traits may be compared in terms of their causal variants to identify shared variants. However, the concept of regional colocalisation can be approached in other ways in the multiple causal variant scenario. One approach reduces the possible hypotheses to two, with the alternative hypothesis corresponding to the existence of a causal variant in a region shared by two (or more) traits.[13] Another focuses on a variant-level definition of colocalisation, estimating the probability that each variant in turn is causal for two traits, whilst allowing that other causal variants (shared or non-shared) may exist in the vicinity[14]. In contrast, the approach proposed here allows the number hypotheses tested to be determined by the data. Whilst it relaxes the assumption of a single causal variant, one obvious caveat is that we have not yet reached (nor may we ever reach) sample sizes which enable all causal variants to be identified. Missed causal variants will provide incomplete comparisons of traits. It is also established that in lower power situations, even Bayesian fine-mapping methods that simultaneously model causal variants may identify a single SNP which tags two or more causal variants[4] and the interpretation of non-colocalisation at such false signals is likely to be misleading. On the other hand, it does seem useful to go beyond asking whether at least one causal variant is shared, and the attempt to both isolate and count the distinct causal variants per trait may be useful in designing follow-up experiments. As we better understand the architecture of complex traits, and design methods that accomodate the multiple causal variants that have been discovered, it is important to bear in mind that results will continue to be limited by sample size, and limited ability to detect rarer variants or those in regions of particular allelic heterogeneity, which even sophisticated methods such as SuSiE may find challenging.

## Availability

Code to perform the simulations may be found at https://github.com/chr1swallace/coloc-susie-paper.

A version of coloc including SuSiE is at https://github.com/chr1swallace/coloc.

## Acknowledgements

We thank Stasia Grinberg and Anna Hutchinson for comments on an earlier version of this manuscript, and Matthew Stephens for detailed explanation of the computational complexities in the `susie_rss` function.

CW is funded by the Wellcome Trust (WT107881) and the MRC (MC UU 00002/4) and supported by the NIHR Cambridge BRC (BRC-1215-20014). The views expressed are those of the author(s) and not necessarily those of the NHS, the NIHR or the Department of Health and Social Care.

This research was funded in whole, or in part, by the Wellcome Trust [WT107881]. For the purpose of Open Access, the author has applied a CC BY public copyright licence to any Author Accepted Manuscript version arising from this submission.

## Supporting Information Captions

**Supplementary Table 1** Results of colocalisation simulations. The columns shown are

- scenario: the simulated causal variants in traits 1 and 2, for example A-AB indicates trait 1 has causal variant A and trait 2 has causal variants A and B
- nsnps_in_region: Number of SNPs in simulated region (1000, 2000, 3000)
- method: method used for coloc analysis. In the case of SuSiE it is appended by the Z treshold used for data trimming
- inferred_cv_pair estimated pair of causal variants under test
- H0,H1,H2,H3,H4 average posterior support for each hypothesis. This is calculated as the sum of posterior probabilities for each hypothesis / number of simulations run. As some variant pairs are unlikely to be tested (eg the pair AA is unlikely to be tested in the scenario A-B) this is not the expected posterior support given AA is tested.

**Supplementary Fig 1.**
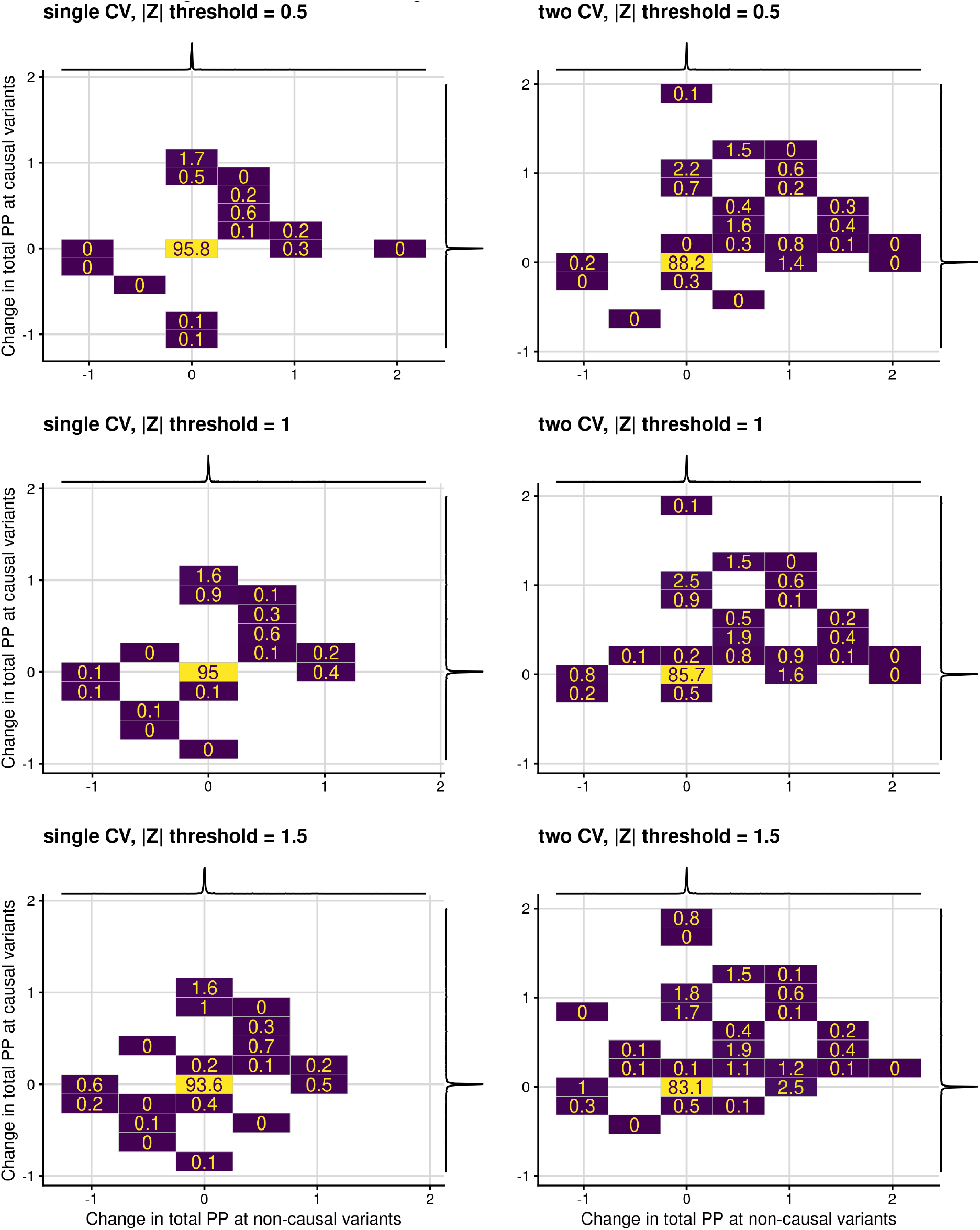
Companion to Fig 1, showing the results for simulated datasets with 3000 SNPs. Legend otherwise as for Fig 1.

**Supplementary Fig 2.**
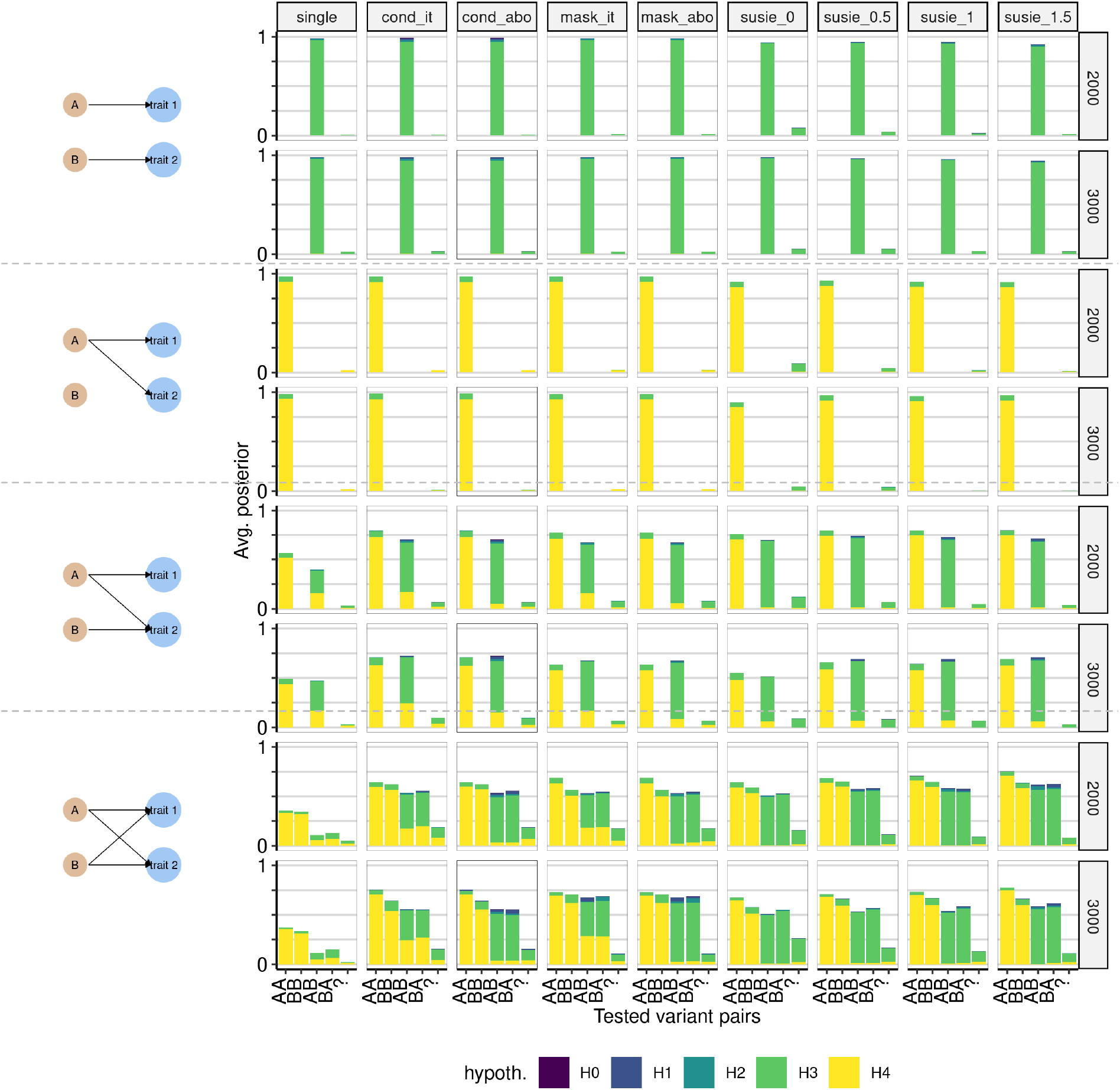
Companion to Fig 3, showing the results for simulated datasets with 2000 or 3000 SNPs. Legend otherwise as for Fig 3.

**Supplementary Fig 3.**
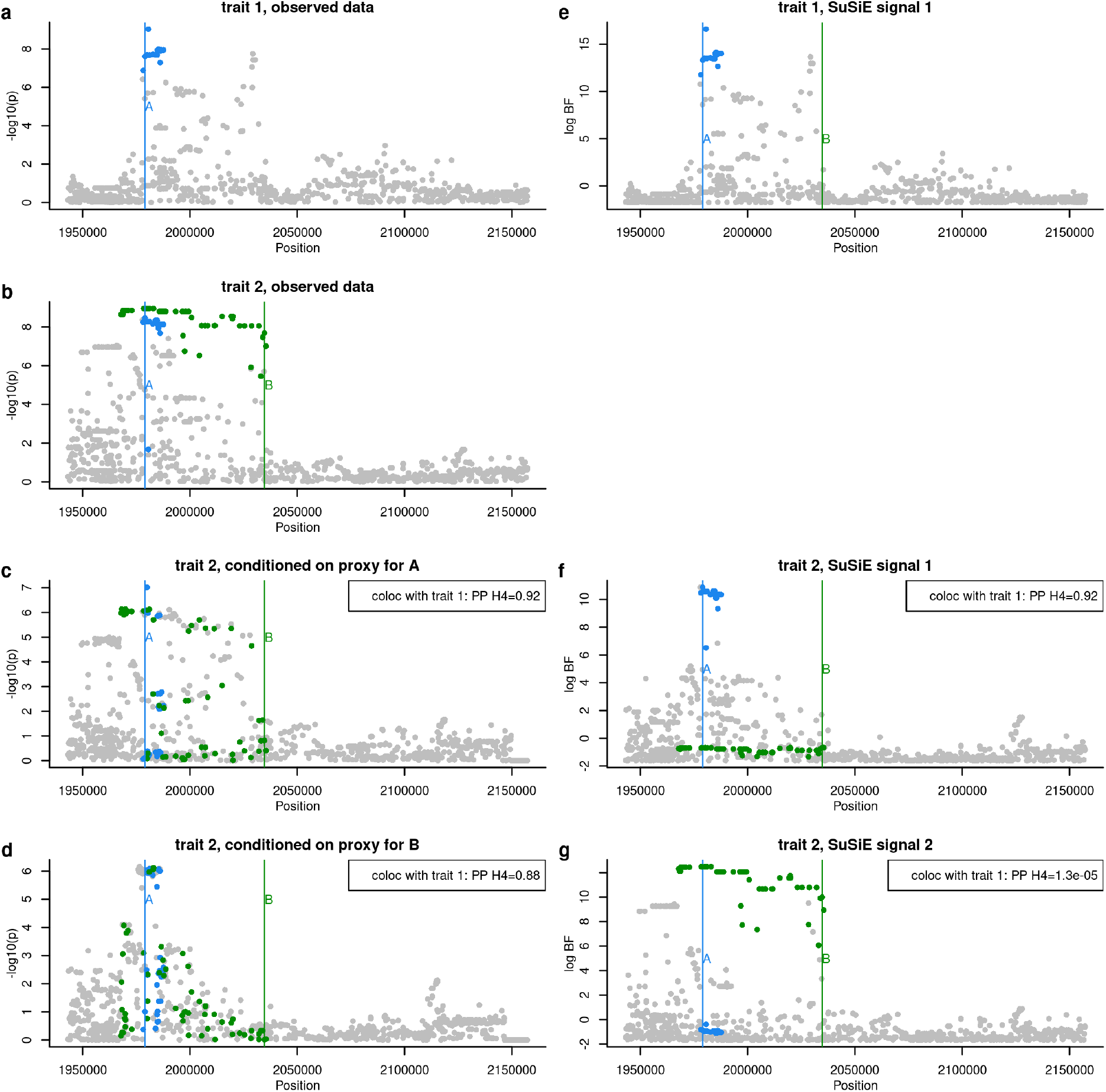
Example where the conditional coloc approach, run in “all but one” mode finds misleading results. **a** and **b** show the observed data (-log p values) for traits 1 and 2 respectively. Conditioning identifies two independent signals for trait 2, and the results of conditioning on the signal closest to causal variants A and B are shown in **c** and **d** respectively. Coloc comparisons are based on (*a, c*) and then (*a, d*). SuSiE analysis of the same data finds one signal in trait 1, and log Bayes factors (BF) for this signal are shown in **e**. It finds two signals for trait 2, and the log BF for these are shown in **f** and **g**. Coloc comparisons are based on (*e, f*) and (*e, g*). The boxes on the lower plots show the results of running coloc analysis on that dataset against the data for trait 1 shown in **a** or **e** as appropriate.

